# Nitrogenous Compound Utilization and Production of Volatile Organic Compounds among Commercial Wine Yeasts Highlight Strain-Specific Metabolic Diversity

**DOI:** 10.1101/2021.03.29.437628

**Authors:** William T. Scott, Oscar van Mastrigt, David E. Block, Richard A. Notebaart, Eddy J. Smid

## Abstract

Strain and environmental nutrient concentrations can affect the production of sensory impact compounds during yeast fermentation. Despite reports on the impact of nutrient conditions on kinetics of cellular growth, it is uncertain to what extent nitrogen utilization by commercial *Saccharomyces cerevisiae* wine strains affects the production of volatile organic (aroma) compounds (VOCs). Here we ask whether i) consumption of amino acids contribute to VOCs (fusel alcohols, acetate esters, and fatty acid esters) in commercial *S. cerevisiae* yeast strains, ii) there is inter-strain variation in VOC production, and iii) there is a correlation between the production of aroma compounds and nitrogen utilization. We analyzed the consumption of nutrients as well as the production of major VOCs during fermentation of a chemically defined grape juice medium with four commercial *S. cerevisiae* yeast strains: Elixir, Opale, R2, and Uvaferm. The production of VOCs was variable among the strains where Uvaferm correlated with ethyl acetate and ethyl hexanoate production, R2 negatively correlated with the acetate esters, and Opale positively correlated with fusel alcohols. The four strains’ total biomass formation was similar, pointing to metabolic differences in the utilization of nutrients to form secondary metabolites such as VOCs. To understand the strain-dependent differences in VOC production, partial least-squares linear regression coupled with genome-scale metabolic modeling was performed with the objective to correlate nitrogen utilization with fermentation biomass and volatile formation. Total aroma production was found to be a strong function of nitrogen utilization (R^2^ = 0.87). We found that glycine, tyrosine, leucine, and lysine utilization were positively correlated with fusel alcohols and acetate esters concentrations e.g., 2-phenyl acetate during wine fermentation. Parsimonious flux balance analysis and flux enrichment analysis confirmed the usage of these nitrogen utilization pathways based on the strains’ VOC production phenotype.

**IMPORTANCE:** *Saccharomyces cerevisiae* is widely used in grape juice fermentation to produce wines. Along with the genetic background, the nitrogen in the environment in which *S. cerevisiae* grows impacts its regulation of metabolism. Also, commercial *S. cerevisiae* strains exhibit immense diversity in their formation of aromas, and a desirable aroma bouquet is an essential characteristic for wines. Since nitrogen affects aroma formation in wines, it is essential to know the extent of this connection and how it leads to strain-dependent aroma profiles in wines. We evaluated the differences in the production of key aroma compounds among four commercial wine strains. Moreover, we analyzed the role of nitrogen utilization on the formation of various aroma compounds. This work illustrates the unique aroma producing differences among industrial yeast strains and suggests more intricate, nitrogen associated routes influencing those aroma producing differences.

## INTRODUCTION

It has been widely recognized that yeast cell growth and overall wine fermentation performance are regulated by initial nitrogen levels within the grape must. Consequently, nitrogen limitation can induce sluggish or stuck fermentations (1–4). Ammonium and amino acids are the primary nitrogen sources used by yeast for general biosynthetic purposes by transferring the amine functional group (5, 6). Not only are yeast cell growth and fermentation completion influenced by the quality and amount of ammonia and amino acids in the grape must, also the production of many crucial volatile organic compounds (VOCs) that are associated with desirable wine bouquet are impacted (7–11). More specifically, these desirable VOCs are higher alcohols and their associated esters and fatty acids. The higher alcohols are products of the Ehrlich pathway, which use branched-chain and aromatic amino acids as substrates (12).

It has been shown that an inverse correlation exists between initial nitrogen levels (excluding at low initial nitrogen levels) and fusel alcohol concentrations (8, 13–16). Furthermore, it has been demonstrated in wine fermentations using *S. cerevisiae* that amino acids are directly involved in the formation of higher alcohols, esters, and fatty acids and these volatile organic compounds subsequently influence aroma attributes of wines (12, 17). Non-volatile compounds, including glycerol, malic acid, and succinic acid, have also been shown to fluctuate depending on nitrogen concentration and source (18–20). Although VOC precursors produced via the Ehrlich pathway have been confirmed, various other amino acids such as alanine, lysine, glycine, histidine, and glutamine could potentially act as precursors or regulators of numerous metabolic pathways linked to aroma compound production. Moreover, surprisingly little is known about the relationship between the dynamics and timing of amino acid utilization and VOC production throughout grape must fermentation. Modulating desirable VOC yield in production strains will allow for valuable process advances in improving wine aroma bouquet, as well as increased flexibility in the production of specific aroma compounds for targeted types of wines. In addition, broader insights into aroma development could potentially lead to the introduction of these qualities into production strains with other desirable characteristics e.g. creating full-bodied aromas in low alcohol wines. Acetate esters and medium-chain fatty acid (MCFA) esters, exhibit a more intricate relationship with initial nitrogen levels because of their biosynthetic routes of production. However, previous work has shown ethyl acetate is positively correlated to medium nitrogen levels (7, 8, 15, 21, 22). A commonly used practice in winemaking is to add nitrogenous compounds to avoid problem fermentations empirically. Although this heuristic is moderately successful, its benefits are inconsistent since the sole addition of inorganic nitrogen as well as improper supplementation of nitrogen have been revealed to negatively impact fermentation performance and aroma compound formation (10, 23). More precisely, low yeast assimilable nitrogen concentrations can cause stuck fermentations and lead to higher H_2_S levels, while high YAN concentrations may cause greater turbidity, stimulate microbial instability and facilitate the formation of unpleasant aromas (24–26). Thus, it would be most advantageous to be able to pre-determine nitrogen levels and the timing of additions to a wine medium to achieve proper aroma character and wine styles, though this is not yet possible with current knowledge of the system.

The numerous *S. cerevisiae* strains, which are selected for winemaking, differ immensely in their aroma production profiles (27–29). For example, a study that used two strains demonstrated the strain with the higher nitrogen requirement formed the higher ester concentration during fermentation of a Chardonnay must (30). Strain-dependent VOC production profiles could be due to variations in how yeast cells metabolize and utilize nitrogenous compounds. Another study, in which three yeast strains were examined in chemically defined media with different nitrogen compositions, indicated measurable differences in volatile and non-volatile compounds especially for the total amount of esters (31). Moreover, Miller et al. (29) showed strain-specific differences in the production of volatile esters in Chardonnay grape juice with varying initial nitrogen levels. Although these studies were groundbreaking in illustrating strain-dependent behavior for several yeast strains under various environmental conditions, they lacked insight regarding the roles specific nitrogenous compounds play in the formation of VOCs during fermentation.

Previous studies (29, 31, 32) have examined the dynamics of the relationship between nitrogen utilization of commercial *S. cerevisiae* strains and the production of VOCs e.g., fusel alcohols and acetate esters. However, earlier studies may be incomplete as it has been suggested that other nitrogen sources or nitrogen-involved metabolic pathways play a role in VOC formation in alcoholic fermentations such as beer (32, 33). The four industrial *S. cerevisiae* strains, Elixir, Opale, R2, and Uvaferm investigated in this work exhibit a range of fermentation and aroma producing capabilities. We illustrated strain-specific behavior affecting the production of key wine associated VOCs. In order to further understand the role of nitrogen utilization in the various aroma-producing attributes among these strains, we applied established principles regarding amino acid degradation. We described how VOCs produced during fermentation correlated with nitrogen (ammonia and amino acids) utilization employing partial least squared (PLS) regression. This multivariate statistical technique was used to elucidate high dimensional data sets to determine possible mechanisms of aroma formation and identify candidate nitrogen sources (ammonia/amino acids) that are essential to these mechanisms. Moreover, we established underlying mechanistic linkages between nitrogen utilization and VOC production using genome-scale metabolic modeling.

## RESULTS

### Fermentations

Fermentations were carried out for each of the *S. cerevisiae* strains in triplicate to evaluate nutrient utilization, i.e. ammonia, amino acids, and sugar consumption, as well as VOC (aroma) production capabilities across strains growing in the same enological medium. All fermentations were conducted under atmospheric conditions at a temperature of 20°C and were performed until completion (t = 404 hr). Throughout the fermentation, cell biomass (estimated from OD600 measurements) and °Brix levels were monitored at 11 time points. Biomass growth, sugar utilization, and ethanol formation curves are shown (see Fig. 1A).

**FIG 1.**
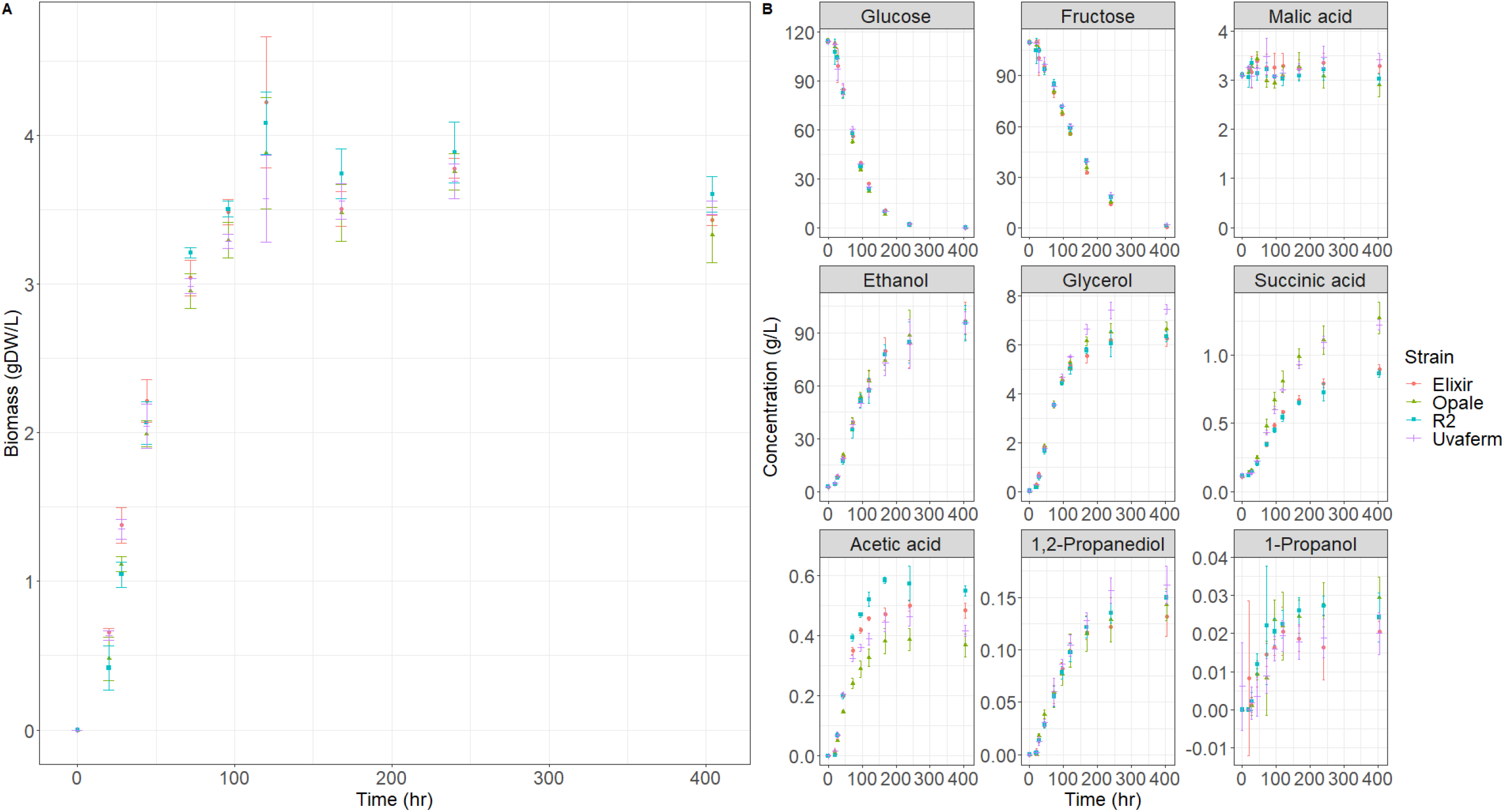
Growth, nutrient consumption, and metabolite production kinetics of the four yeast strains in MMM medium depicting (A) dry cell weight (biomass) and (B) major metabolites over the course of the fermentations. Error bars represent standard deviations (n = 3).

All strains reached maximum biomass at approximately 120 hr. Furthermore, all strains do not significantly differ in biomass concentrations (One-way ANOVA, p = 0.974). All cultures were fermented to completion (less than 4g/L residual sugar (34)). More specifically, fermentations performed with Uvaferm, R2, Opale, and Elixir strains resulted in total final sugar concentrations of 1.6 g/L, 0.9 g/L, 1.3 g/L, and 0.3 g/L, respectively. All the strains reached similar final ethanol concentrations ranging between 90 and 100 g/L. The glycerol, malic acid, acetic acid, lactic acid, succinic acid, and 1,2 propanediol production were measured, as these compounds are vital to the sensory characteristics and stability of wines. Glycerol concentrations found in the cultures ranged from 6.3 g/L for R2 to 7.5 g/L for Uvaferm (see Fig. 1B). These values conform to those found in wine (35). For all strains, malic acid levels were relatively constant throughout the fermentation at approximately 3 g/L (Fig. 1B). Acetic acid production levels varied between the strains with all of the final concentrations in the range of 350 mg/L for Opale to 550 mg/L for R2 (Fig. 1B). Succinic acid concentrations reached by the strains ranged from 0.8 g/L for R2 to 1.3 g/L for Opale (Fig. 1B). Final concentrations of 1,2 propanediol were relatively similar among the strains varying from 109 mg/L for Elixir to 160 mg/L for Uvaferm (Fig. 1B). One-way ANOVA revealed (95% confidence interval) that the production of acetic acid, malic acid, and succinic acid was significantly different among all of the strains (p < 0.05). In contrast, concentrations of the other six major metabolites (Fig. 1B) did not differ significantly among the strains (p > 0.05).

### Nitrogenous compound utilization and VOC production profiles during fermentation

The concentrations of ammonia and amino acids were measured over the course of the fermentations (Fig. 2). Amino acid and ammonia consumption profiles deviated between all of the strains illustrating strain-dependent behavior for some compounds such as alanine, glycine, asparagine, leucine, and phenylalanine. Moreover, concentrations of other amino acids such as serine, valine, lysine, methionine, and threonine, differed significantly between across two or three strains. Overall, the level of consumption for 15 out of 20 amino acids studied differed significantly (ANOVA, p < 0.05). However, these and most variations in amino acid concentrations were observed at 20 and 28 hours. As noticed (Fig. 2), it is evident that ammonia is consumed most rapidly, even before amino acids in the synthetic must. All of the amino acids were consumed during the fermentations except proline, which was not taken up and remained in the medium until the end of fermentation (data not shown).

**FIG 2.**
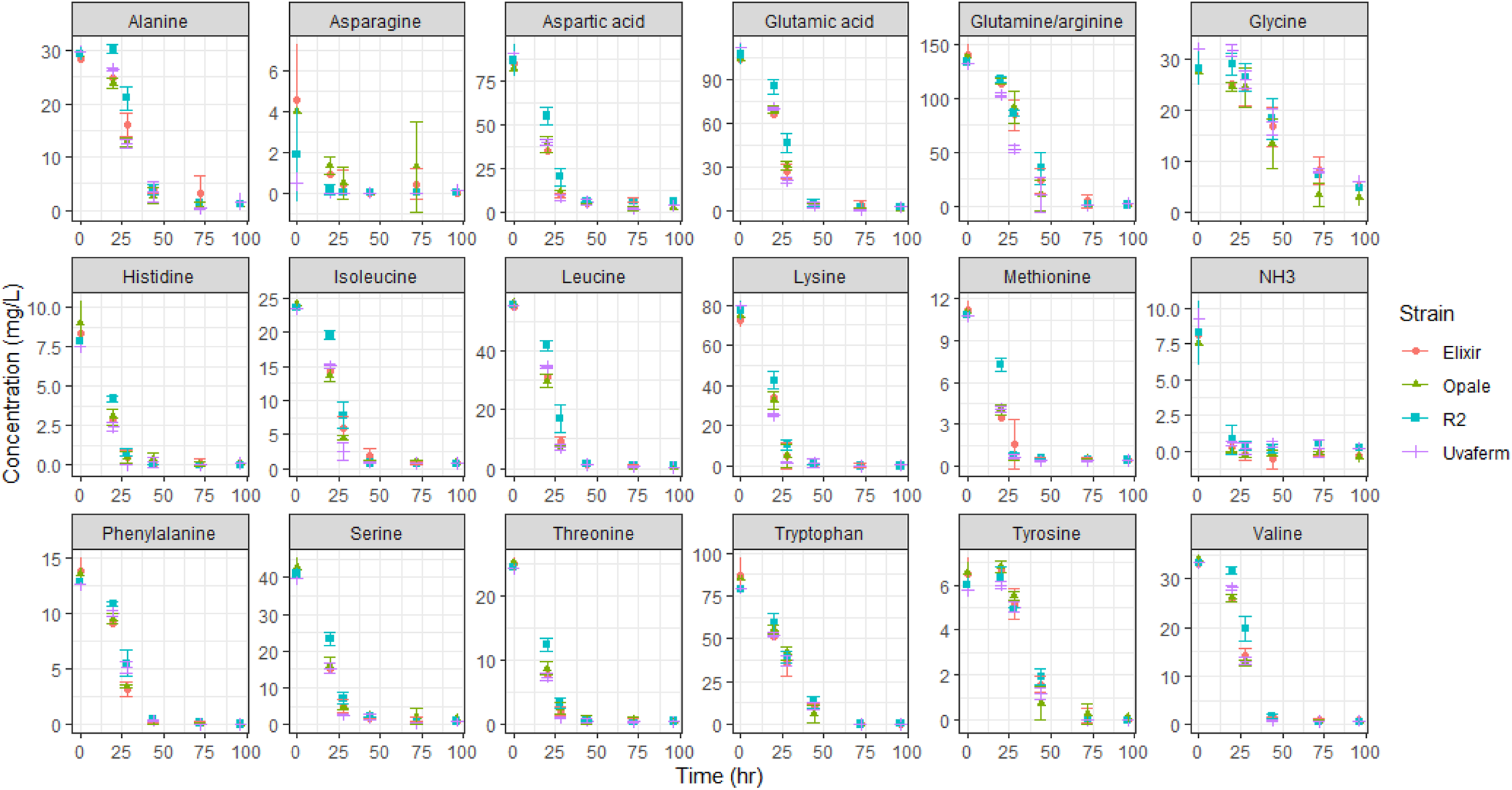
Amino acid and ammonia consumption of the four yeast strains in MMM medium showing the change of amino acid concentrations during fermentation. Error bars represent standard deviations (n = 3).

Many amino acids were mainly consumed during the exponential cell growth stage of fermentation. Only some small amounts of Ala, Asn, Gly, and Met, remained in the media after 100 hours, though all are utilized shortly after that. The consumption data reveal preferences of yeast for particular amino acids during utilization over the exponential growth phase. The consumption of the nitrogenous compounds can be separated into four groups based on the time at which 95% utilization occurs from the compound. Group I consists of earliest consumed compounds where strain-dependent consumption NH_3_, Asn, Lys, Met, Ile (Uvaferm only), and Thr (Uvaferm only) occurs with 95% of their concentrations being drastically depleted by 28 hours. Group II contains subsequently preferred compounds where 95% of the compound was consumed by 44 elapsed hours. These compounds are Arg, Gln, Ile, Leu, Phe, Ser, Thr, His, Asp, and Val. Group III consists of compounds such as Ala, Trp, and Tyr which all show steady consumption until 95% of their concentrations were utilized by 72 hours. A remaining group, Group IV, was taken up after some initial delay (not consumed within the first 20 hours) and did not have 95% utilization until after 96 hours. This sole amino acid was Gly.

To study the relationship between amino acid consumption and VOC formation during grape juice fermentation, 10 VOCs (aromas) were measured using HS SPME GC-MS from various compound classes pertaining to different aroma properties throughout the fermentation process (Fig. 3). The compounds consisted of four fusel alcohols: isobutanol, isoamyl alcohol, 2-phenyl ethanol, and methionol; four acetate esters: ethyl acetate, isobutyl acetate, isoamyl acetate, and 2-phenyl ethyl acetate; and two ethyl esters: ethyl butanoate and ethyl hexanoate. The kinetics of the measured VOCs formation was dependent on the yeast strain. In addition, the maximum and final concentration of VOCs varied significantly among the strains (ANOVA, p < 0.05). In general, the VOC concentrations of all of the strains significantly differed by 168 hours of fermentation. The only compounds that did not differ significantly at the end of fermentation were 2-methyl-1-propanol and 2-methyl-1-propyl acetate.

**FIG 3.**
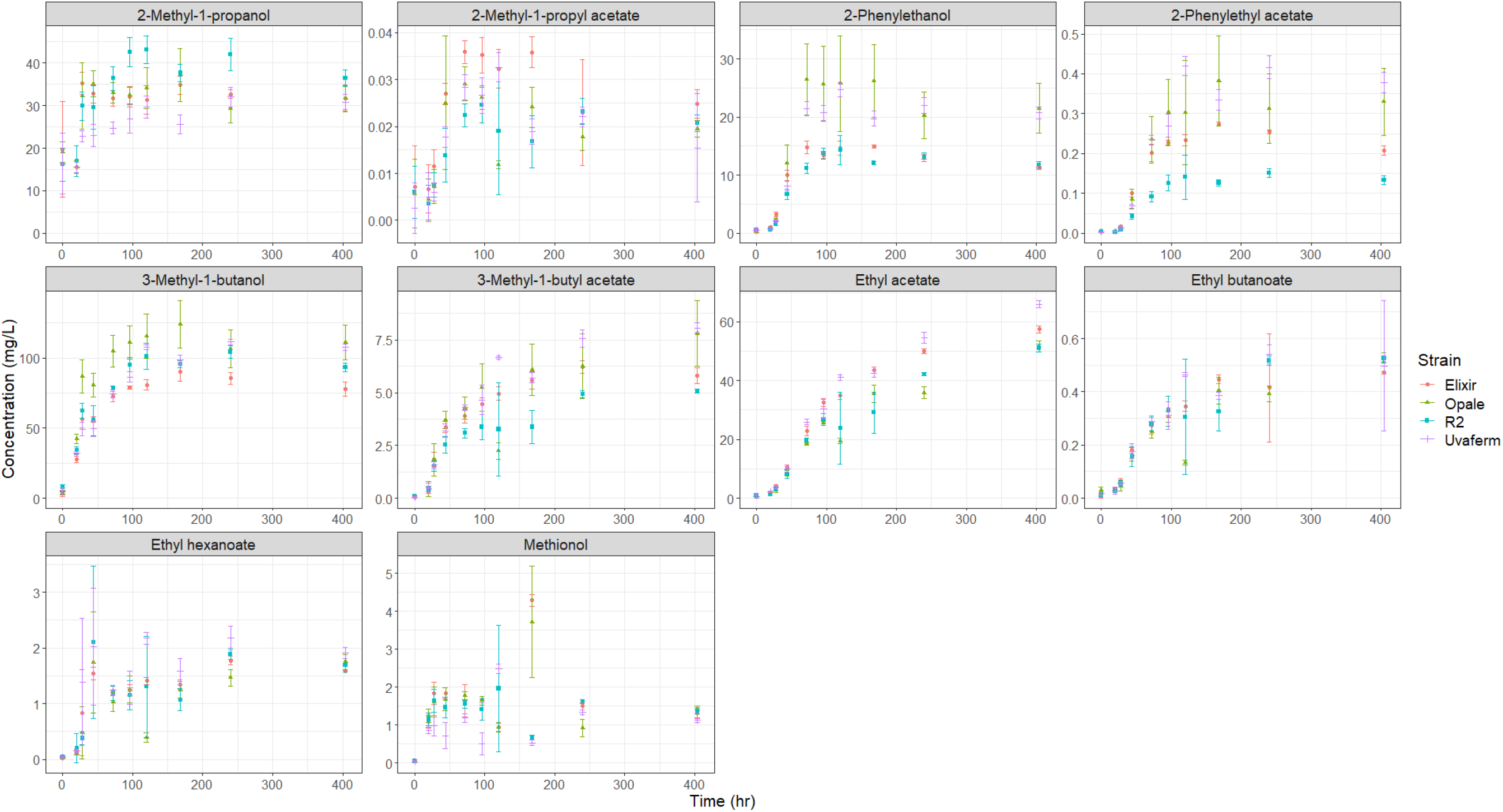
VOC (aroma) production of the four yeast strains in MMM medium showing the change of VOC concentrations during over the course of the fermentations. Standard deviations are represented (n = 3).

All VOCs begin to form early in the fermentation process. Some compounds such as 3-methyl 1-butanol and methionol were even generated before 20 hours though nearly all VOCs formation characteristically corresponds with yeast growth. Moreover, the maximum production rate of fusel alcohols and acetate esters occurred from 28 hours to 96 hours. Fusel alcohols were produced at the highest concentrations with isoamyl alcohol, isobutanol, and 1-propanol showing the most rapid production earliest during fermentation. The acetate ester production rate began to decline for all of the strains after 96 hours and became relatively stagnant after 168 hours. Profiles for ethyl esters were similar to those observed for acetate esters. However, initial ethyl ester formation proceeded at a lower rate than acetate esters, and the maximum concentrations were much less than acetate esters. Overall, most VOCs stagnated production after 96 hours, which indicates growth-dependent behavior. Some VOCs, in particular, 2-methyl-1-propyl acetate, show a decrease in concentration towards the end of fermentation most likely due to volatilization. All final VOCs concentrations were found within the typical range found in wines (28).

In order to compare the results of volatile production by the strains, a principal component analysis (PCA) of the VOCs was performed. From the PCA, 76.26% of the variance was explained by the first two principal components (PC) (PC1 = 51.25% and PC2 = 24.98%). As depicted, separation of the samples was achieved according to the yeast strains (Fig. 4). PC1 separated R2 from the other three strains, while PC2 separated Elixir, Opale and Uvaferm strains. Although Lallemand characterizes Uvaferm as a neutral aroma-imbuing strain, loadings for ethyl hexanoate and ethyl acetate were correlated with the Uvaferm strain indicating relatively higher production for Uvaferm. Opale and R2 are described as conveying citrus, fruity, and floral aromas to wines from enhanced ester production and low H_2_S and SO_2_ formation. Nevertheless, higher loadings for fusel alcohols were correlated with the Opale strain along PC2, whereas loadings for the acetate esters were negatively correlated with the R2 strain indicating relatively low acetate ester formation. The manufacturer claims that Elixir produces a wide array of beneficial fatty acid esters while limiting formation of acetate esters. As depicted, it appears Elixir has a correlation with ethyl butanoate and ethyl acetate (Fig. 4).

**FIG 4.**
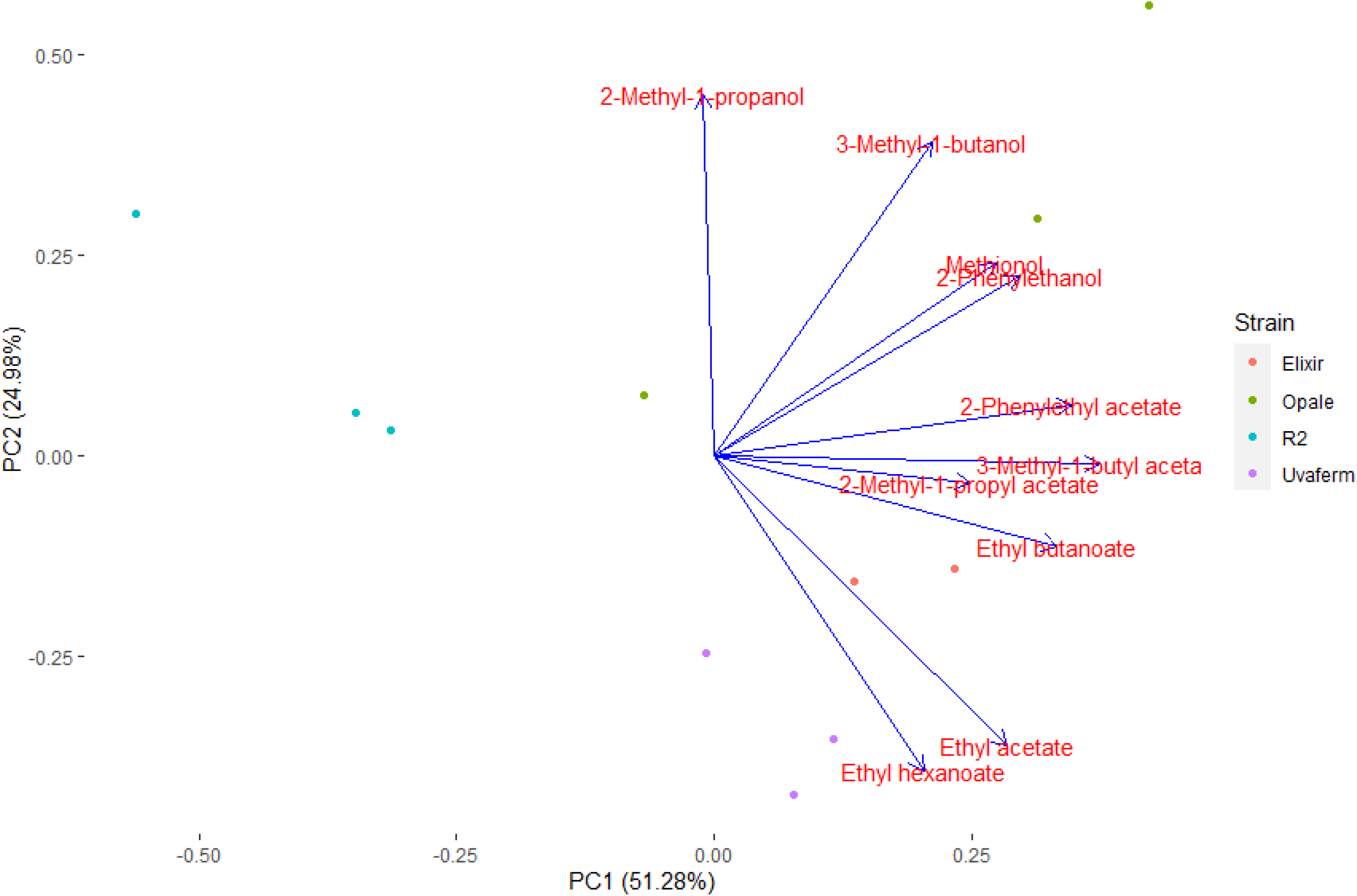
PCA scores and loadings plot of PC1 and PC2 derived from the volatile compounds produced by the yeast strains at t = 168 hours.

### Correlations of nitrogenous compound utilization with VOC formation

One of the main hypotheses of this study asserted that utilization of ammonia and amino acids during fermentation contributed to the characteristic difference in the formation of VOCs among the yeast strains. Firstly, ammonia and amino acids serve as an essential nitrogen source promoting good growth of the *S. cerevisiae* culture. Therefore, enhancing growth of yeast cultures leads to the overall increase of production of VOCs and VOC precursors. Secondly, it is known that specific amino acids are utilized and degraded which contribute to the synthesis, via the Ehrlich pathway, of fusel alcohols and subsequently to the production of acetate esters. However, other pathways related to lysine and glycine degradation, which have been shown to play a role in VOC formation in other types of fermentations may be essential to enological fermentations as well. To evaluate this hypothesis and understand the causes of strain-dependent variation, the ammonia and amino acid utilization for all four yeast strains were measured and correlated with biomass formation and VOC production for 11 different aroma compounds during exponential growth phase (Table 1).

**TABLE 1.** Summary of results from partial least-squares regression analysis

| **Response variable**<br>**(predictor variable)** | **No. of**<br>**latent**<br>**variables** | **RMSEC** | **RMSECV** | **R<sup>2</sup>** | **Q<sup>2</sup>** | **%**<br>**variance**<br>**captured**<br>**X-block** | **%**<br>**variance**<br>**captured**<br>**Y-block** |
| --- | --- | --- | --- | --- | --- | --- | --- |
| Biomass concn.<br>(nitrogen util.) | 10 | 0.24 | 0.30 | 0.96 | 0.93 | 100.00 | 95.52 |
| Fusel alcohol concn.<br>(nitrogen util.) | 5 | 47.25 | 53.11 | 0.87 | 0.83 | 100.00 | 87.02 |
| 1-Propanol concn.<br>(nitrogen util.) | 9 | 5.35 | 6.92 | 0.92 | 0.87 | 100.00 | 92.02 |
| 3 Methyl-1-butanol<br>concn. (nitrogen util.) | 7 | 35.56 | 40.57 | 0.82 | 0.77 | 100.00 | 82.32 |
| 2 Methyl-1-propanol<br>concn. (nitrogen util.) | 8 | 7.69 | 9.44 | 0.80 | 0.71 | 99.79 | 80.17 |
| 2 Phenylethanol concn.<br>(nitrogen util.) | 4 | 7.09 | 7.68 | 0.85 | 0.82 | 100.00 | 84.82 |
| Methionol concn.<br>(nitrogen util.) | 8 | 0.64 | 0.73 | 0.61 | 0.50 | 100.00 | 61.01 |
| Acetate ester concn.<br>(nitrogen util.) | 12 | 12.83 | 16.69 | 0.84 | 0.74 | 99.95 | 84.16 |
| Ethyl acetate concn.<br>(nitrogen util.) | 11 | 11.10 | 14.85 | 0.85 | 0.73 | 99.97 | 84.48 |
| 3 Methyl-1-butyl<br>acetate concn. (nitrogen<br>util.) | 12 | 1.51 | 2.06 | 0.85 | 0.74 | 99.91 | 85.43 |
| 2 Methyl-1-propyl<br>acetate concn. (nitrogen<br>util.) | 14 | 0.01 | 0.01 | 0.85 | 0.74 | 100.00 | 85.42 |
| 2 Phenylethyl acetate<br>concn. (nitrogen util.) | 12 | 0.10 | 0.13 | 0.82 | 0.72 | 99.97 | 82.30 |
| Fatty acid ethyl ester<br>concn. (nitrogen util.) | 4 | 0.73 | 0.81 | 0.77 | 0.72 | 100.00 | 77.19 |
| Ethyl butanoate concn.<br>(nitrogen util.) | 11 | 0.12 | 0.16 | 0.86 | 0.75 | 99.98 | 86.09 |
| Ethyl hexanoate concn.<br>(nitrogen util.) | 5 | 0.67 | 0.70 | 0.77 | 0.71 | 100.00 | 77.31 |
| Total aroma concn.<br>(nitrogen util.) | 5 | 58.62 | 65.73 | 0.87 | 0.84 | 100.00 | 87.08 |

\*nitrogen util. – ammonia and amino acid utilization

In order to gain metabolic insight into how nitrogen utilization impacts characteristic VOC formation among the strains, the degree of the correlation between nitrogen consumption and VOC formation was evaluated using multivariate statistics. Since the datasets are high-dimensional, PLS linear regression was employed to correlate the contributions of ammonia and individual amino acid utilization to the production of biomass and VOCs by the yeast strains during fermentation. Statistical determination of the nitrogenous variables that contribute most to the variation in the fermentation kinetic data regarding VOCs was performed using interval-partial least squared (iPLS) regression analysis. iPLS was done using the ammonia and amino acid utilization of all four *S. cerevisiae* strains during the initial five time points covering the near complete uptake of the nitrogen during the first 96 hours of fermentation. The biomass dry cell weight concentration, individual VOC concentration, each VOC class (i.e. fusel alcohol concentration, acetate ester concentration, and fatty acid ethyl ester concentration), and total VOC concentration were the response variables that comprised the Y-block in 16 different PLS regressions. The response variables contained concentration measurements corresponding to the same time points as the predictor variables. From the initial duration of the fermentation until 96 hours, the ammonia and amino acid utilization data were input as the predictor variables used to develop the X-block in each of the 16 PLS regressions. Subsequently, the PLS regression analysis determined which variables in both the Y-block and the X-block contributed most to the variation in the data. Lastly, the PLS regression analysis indicated how the variables were correlated and then lumped the variables into a new latent variable (LV). Our summarized results show that nitrogen utilization is strongly correlated with biomass formation, and production of each of the aroma compounds except methionol for which only moderate correlation was found (Table 1). The reason could be due to methionol production being controlled by sulfur uptake.

PLS regression was first conducted to determine how nitrogen utilization contributed to yeast growth (biomass formation) during the exponential growth phase in the fermentation. Table 1 summarizes the results from this analysis, where it lists that this model generated 10 LVs encompassing 100% of the variation in the nitrogen utilization data and 95.5 in the biomass concentration data. The PLS regression analysis yielded a correlation coefficient of R^2^ = 0.96, which signified a robust linear relationship between the measured biomass dry cell concentration versus the predicted biomass dry cell concentration according to the nutrient utilization of these yeast strains (Fig. S1). Cross-validation (CV) was performed in order to prevent the model from overfitting the data (i.e. model being applicable for the test observations and not being applicable for new observations) and to evaluate how the model would operate using a new set of data. CV correlation coefficients (Q^2^) were generated to illustrate the predictive strength of the model (36). The root mean squared error of correlation (RMSEC) for the biomass concentration as a function of nitrogen utilization was 0.24 gDW/L of biomass, the root mean squared error in cross-validation (RMSECV) was 0.30 gDW/L of biomass, and the Q^2^ value for this model was 0.93, which highlights that nitrogen utilization was an excellent predictor of the biomass concentration reached by these strains (Fig. S1).

A separate series of PLS regression analyses were performed subsequently for each of the VOCs as well as classes of compounds to investigate a correlation between the production of VOCs during the fermentations and nitrogen utilization of the yeast strains. Each of the models captured at least 99.5% of the variation in the nitrogen utilization data and at least 77% of the variation in the aroma concentration data was captured by each of the models except methionol (Table 1). In addition, the RMSEC and RMSECV for the individual aroma compounds and aroma compound classes as a function of ammonia and amino acid utilization are summarized in Table 1. The R^2^ value for each model for VOCs achieved an R^2^ value greater than 0.77 indicating nitrogen utilization was correlated with the production of each VOC. The only aroma compound that the PLS regression analysis showed to have a more modest correlation with nitrogen utilization was methionol. Partial least squares regression modeling yielded eight LVs and generated a coefficient of determination (R^2^) of 0.61. The model cross-validation results indicate a Q^2^ of 0.5 and RMSEC and RMSECV values of 0.64 mg/L and 0.73 mg/L of methionol, respectively. Overall, these data point to a modest correlation between the methionol concentration and nitrogen utilization during fermentation. As a result, it is suggested that not only nitrogen utilization determines the production of methionol during fermentation and that there may be other processes that play a role in methionol production such as sulfur utilization.

### Specific nitrogenous compounds associated with the formation of VOCs

One of the core goals of this work was to improve understanding of the impacts of nitrogen utilization on VOC profile differences among four commercial wine yeast strains. By applying PLS to the datasets mentioned earlier, information about how specific nitrogen utilization variables correlate with each of the 16 response variable sets was obtained. This facilitated understanding which nitrogenous compounds are responsible for aroma production and might offer clues about metabolic variations among the strains. The PLS models were able to explain the biomass formation and the production of each of the aroma compounds, excluding methionol, which was moderately predicted. Regression vector plots were created to assess the statistical weights of the original nitrogen predictor variables on the PLS models. In other words, to determine the degree to which the 18 original nitrogen variables were positively or negatively correlated with the response variable. These regression vector plots for PLS models predicting individual aroma compounds and classes of aroma compounds are provided in the supplementary material (Fig. S2), while a summary is provided in Table 2. In general, iPLS selection yielded more variables for ester models than for fusel alcohol models. All models shared many similar nitrogenous variables; however, most nitrogenous variable combinations were unique to each model for VOC predictions (Table 2).

**TABLE 2.**
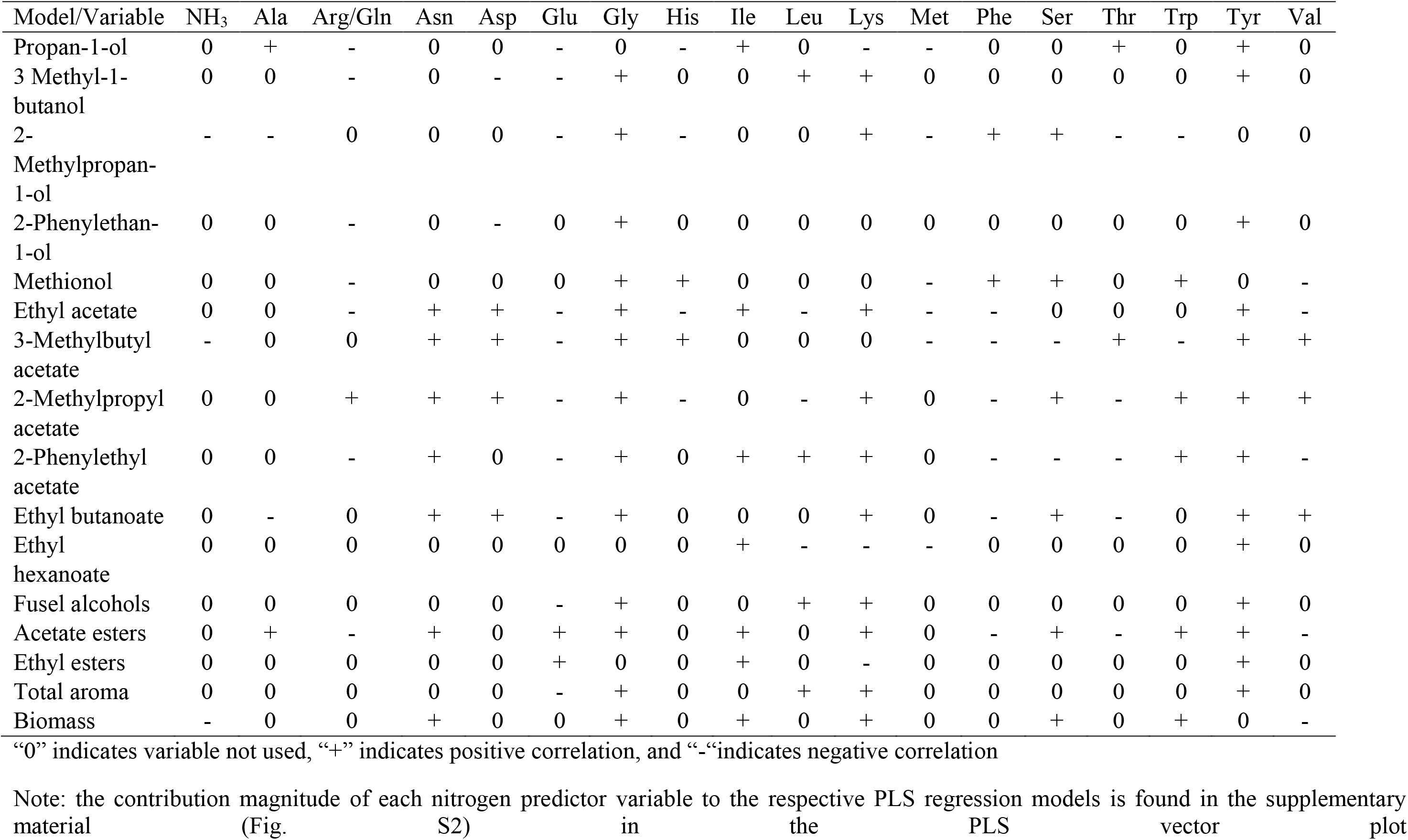
Summary of results from partial least-squares regression analysis vector plots of the predictor variables

| Model/Variable | NH<sub>3</sub> | Ala | Arg/Gln | Asn | Asp | Glu | Gly | His | Ile | Leu | Lys | Met | Phe | Ser | Thr | Trp | Tyr | Val |
| --- | --- | --- | --- | --- | --- | --- | --- | --- | --- | --- | --- | --- | --- | --- | --- | --- | --- | --- |
| Propan-1-ol | 0 | + | - | 0 | 0 | - | 0 | - | + | 0 | - | - | 0 | 0 | + | 0 | + | 0 |
| 3 Methyl-1-butanol | 0 | 0 | - | 0 | - | - | + | 0 | 0 | + | + | 0 | 0 | 0 | 0 | 0 | + | 0 |
| 2-Methylpropan-1-ol | - | - | 0 | 0 | 0 | - | + | - | 0 | 0 | + | - | + | + | - | - | 0 | 0 |
| 2-Phenylethan-1-ol | 0 | 0 | - | 0 | - | 0 | + | 0 | 0 | 0 | 0 | 0 | 0 | 0 | 0 | 0 | + | 0 |
| Methionol | 0 | 0 | - | 0 | 0 | 0 | + | + | 0 | 0 | 0 | - | + | + | 0 | + | 0 | - |
| Ethyl acetate | 0 | 0 | - | + | + | - | + | - | + | - | + | - | - | 0 | 0 | 0 | + | - |
| 3-Methylbutyl acetate | - | 0 | 0 | + | + | - | + | + | 0 | 0 | 0 | - | - | - | + | - | + | + |
| 2-Methylpropyl acetate | 0 | 0 | + | + | + | - | + | - | 0 | - | + | 0 | - | + | - | + | + | + |
| 2-Phenylethyl acetate | 0 | 0 | - | + | 0 | - | + | 0 | + | + | + | 0 | - | - | - | + | + | - |
| Ethyl butanoate | 0 | - | 0 | + | + | - | + | 0 | 0 | 0 | + | 0 | - | + | - | 0 | + | + |
| Ethyl hexanoate | 0 | 0 | 0 | 0 | 0 | 0 | 0 | 0 | + | - | - | - | 0 | 0 | 0 | 0 | + | 0 |
| Fusel alcohols | 0 | 0 | 0 | 0 | 0 | - | + | 0 | 0 | + | + | 0 | 0 | 0 | 0 | 0 | + | 0 |
| Acetate esters | 0 | + | - | + | 0 | + | + | 0 | + | 0 | + | 0 | - | + | - | + | + | - |
| Ethyl esters | 0 | 0 | 0 | 0 | 0 | + | 0 | 0 | + | 0 | - | 0 | 0 | 0 | 0 | 0 | + | 0 |
| Total aroma | 0 | 0 | 0 | 0 | 0 | - | + | 0 | 0 | + | + | 0 | 0 | 0 | 0 | 0 | + | 0 |
| Biomass | - | 0 | 0 | + | 0 | 0 | + | 0 | + | 0 | + | 0 | 0 | + | 0 | + | 0 | - |

“0” indicates variable not used, “+” indicates positive correlation, and “-” indicates negative correlation
Note: the contribution magnitude of each nitrogen predictor variable to the respective PLS regression models is found in the supplementary material (Fig. S2) in the PLS vector plot

The regression vector plots for all the yeast strains illustrate the degree to which the 18 original nitrogen predictor variables are correlated with the response variables (VOC and biomass concentrations) (Fig S2). Furthermore, it is worth noting that the magnitude of the correlation coefficients for each nitrogen predictor variable was different across the PLS regression models (Fig S2). The model for biomass growth indicated that asparagine, serine, glycine, lysine, isoleucine, and tryptophan were all positively correlated with biomass concentration, whereas, histidine, threonine, and valine are negatively correlated with biomass concentration (Table 2). Ammonia was shown to have a very slight negative correlation with biomass growth with a correlation coefficient, less than 0.1, but this contribution could be ascribed to measurement artefacts as more than one type of assay was needed to determine ammonia concentrations. The model for fusel alcohols indicated that Gly, Lys, Tyr, and Leu were positively correlated with fusel alcohol concentration while Glu was negatively correlated with fusel alcohol concentration (Table 2). Correspondingly, the model for acetate esters shared many of the same correlations with predictor variables as the fusel alcohol model. However, Ala, Asp, Ile, Ser, and Trp were also found to be positively correlated with acetate ester concentration, while Val, Arg, Gln, and Phe were negatively correlated with acetate ester concentration. Lastly, the ethyl esters model indicated that Glu, Ile, and Tyr positively correlated with ethyl ester concentration whereas Lys negatively correlated with ethyl ester concentration.

A summary of the results from all the regression vector plots highlights correlative behavior of the nitrogen variables for predicting individual VOC production (Table 2). The regression vector plots revealed some patterned behavior regarding correlation found for particular nitrogenous compounds. Ammonia, methionine, and glutamic acid were only negatively correlated with the prediction of aroma compound production. Furthermore, arginine and glutamine negatively correlated with the prediction of most fusel alcohols and acetate esters except 2-methyl propyl acetate, which was positively correlated. Conversely, some nitrogen variables were only positively correlated with aroma production meaning that a higher consumption of these nitrogen compounds leads to higher aroma concentrations. These nitrogen variables were asparagine, isoleucine, and tyrosine. Glycine positively correlated with all aroma compounds except 1-propanol and ethyl hexanoate, and aspartic acid positively correlated with all acetate esters except 2-phenylethyl acetate. Nitrogen variables that were positive correlated to the total aroma model were glycine, leucine, lysine and tyrosine. Lysine positively correlated with many fusel alcohols and acetate esters. The results for the remaining nitrogen variables were mixed concerning the prediction of aroma compound.

### Parsimonious flux balance analysis and flux enrichment analysis

The above described PLS correlations raise the hypothesis that amino acid utilization pathways are enriched in activity within the entire metabolic network. To test this, we applied parsimonious flux balance analysis (pFBA) and flux enrichment analysis (FEA) with fermentation data used as constraints. The pFBA analysis revealed many essential genes / reactions for our modeled system (see Table 3). Furthermore, many of the essential reactions, as well as non-essential pFBA optimal and less efficient metabolic reactions (ELE and MLE reactions, see Methods for detailed information), are related to amino acid degradation and VOC forming pathways (Fig. 5). The activity of less efficient reactions may point to a trade-off between growth and VOC production, i.e., when VOCs are produced also less efficient reactions must occur in context of growth. After excluding transport reactions and non-enzyme associated reactions, there are 891 essential reactions where 204 essential reactions are associated with amino acid degradation pathways, representing 22.9% of the essential reactions. The number of reactions in each category of amino acid metabolism shown to be participating essential reactions are outlined (Table 3).

**FIG 5.**
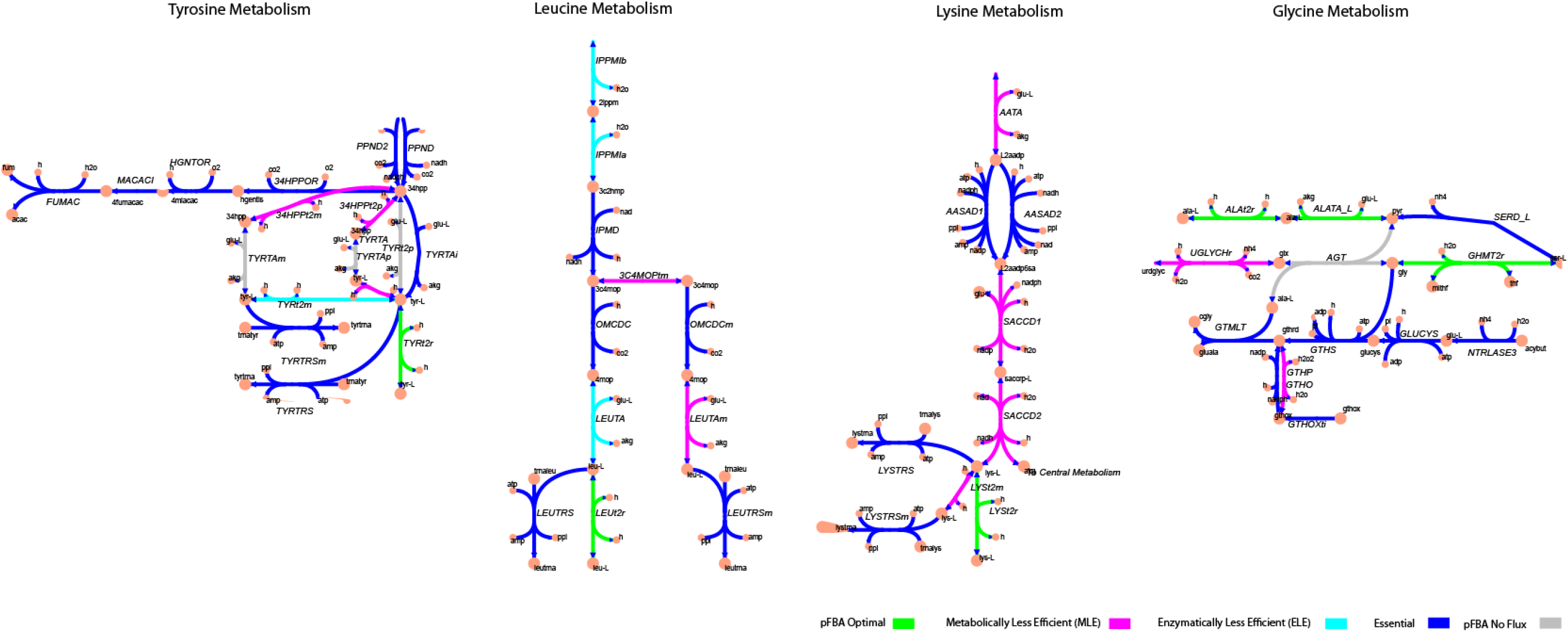
Key *S. cerevisiae* metabolic pathways illustrating reaction classes from applying parsimonious flux balance analysis. All reactions and metabolites shown in the figure contain Biochemical Genetic and Genomic (BiGG) database identifiers (http://bigg.ucsd.edu/).

**TABLE 3.** Summary of the essential reactions and the flux enrichment analysis results of yeast model subsystems

| **GSMM Metabolism Group** | **No. of**<br>**Essential**<br>**Reactions** | **Enriched**<br>**set size** | **p-value** |
| --- | --- | --- | --- |
| Sterol Metabolism | 36 | 49 | 4.80E-31 |
| Tyrosine Tryptophan and<br>Phenylalanine Metabolism | 32 | 44 | 1.23E-26 |
| Complex Alcohol Metabolism | 33 | 41 | 3.13E-24 |
| Arginine and Proline Metabolism | 23 | 33 | 1.79E-18 |
| Methionine Metabolism | 16 | 20 | 1.40E-10 |
| Glycine and Serine Metabolism | 18 | 19 | 4.96E-10 |
| Threonine and Lysine Metabolism | 14 | 19 | 4.96E-10 |
| Valine Leucine and Isoleucine<br>Metabolism | 17 | 19 | 4.96E-10 |
| Glycolysis and Gluconeogenesis | 19 | 22 | 1.06E-09 |
| Pyruvate Metabolism | 17 | 20 | 1.40E-09 |
| Glutamate Metabolism | 12 | 17 | 5.94E-09 |
| Other Amino Acid Metabolism | 5 | 13 | 7.13E-07 |
| Glycerolipid Metabolism | 9 | 12 | 2.28E-06 |
| Cysteine Metabolism | 10 | 10 | 2.24E-05 |
| Histidine Metabolism | 10 | 11 | 5.89E-05 |
| Alanine and Aspartate Metabolism | 8 | 9 | 6.88E-05 |
| Phospholipid Metabolism | 7 | 8 | 0.00021 |
| Fructose and Mannose Metabolism | 5 | 6 | 0.00186 |
| Citric Acid Cycle | 10 | 10 | 0.00198 |
| Glutamine Metabolism | 4 | 4 | 0.01578 |

After obtaining the essential reactions, we employed flux enrichment analysis (FEA) examining all of the metabolic to identify statistically significantly enriched reactions within the subsystems. We hypothesized that nitrogenous compounds (i.e., amino acids, ammonium, etc.) play an appreciable role in VOC formation throughout enological fermentation. Here, we see that from flux enrichment analysis (FEA) of the essential reactions determined by pFBA, delivers a classification of the metabolic groups within the GSMM. A ranking where a position in the list of analyzed pathways is arranged by significance is given (Table 3) (37). Moreover, the FEA highlights tyrosine, glycine, lysine, and leucine metabolism and complex alcohol metabolism (fusel alcohols) as the most significant parts of metabolism when constrained with our experimental data. Thus, FEA supported the results from our PLS regression analysis and together with pFBA it highlighted which reactions are essential. We also found that sterol (lipid metabolism), arginine, and methionine (sulfur) metabolism play significant roles. This result could indicate substantial influence coming from lipid metabolism and sulfur metabolism. Despite potential influence from other parts of metabolism on VOC formation, our statistical and metabolic modeling results firmly indicated that amino acid utilization routes such as leucine, glycine, lysine, and tyrosine induce VOC production (Fig. 5). We also observed within these amino acid utilization routes the essential reactions and less efficient routes (MLE and ELE) (Fig. 5). Interestingly, it was observed that transaminase (e.g., TYRTAi) and decarboxylase associated reactions (e.g., OMCDC) were essential, indicating that Ehrlich or amino acid degradation pathways are involved in VOC formation. Nevertheless, other reactions were also found to be essential such as reactions associated with acetate (e.g., FUMAC) suggesting to other routes influence VOC formation as well.

## DISCUSSION

Fermentations were performed with four commercial *S. cerevisiae* strains with diverse aroma bouquet producing potential and growth characteristics under identical experimental conditions to analyze the nitrogenous compounds that correlated with yeast cell growth and aroma production. In agreement with previous studies, these experimental results illustrate significant differences in volatile and non-volatile production depending on the applied yeast strain (10, 29, 38, 39). In most cases, the fermentations performed with the Opale strain had the highest maximum and final non-volatile and volatile concentrations. In contrast, the fermentations conducted using the R2 strains had the lowest maximum and final non-volatile and volatile concentrations. Although once normalizing the non-volatile and volatile concentrations to the biomass formed, the differences among the strains are less evident (data not shown). Furthermore, when examining the product specifications from Lallemand (Lallemand, Montreal, Quebec), it is presumed that the Uvaferm strain, which is a neutral aroma producer, would generate the least volatile compounds under the same medium conditions relative to the other strains. Our results suggest, as shown in previous studies, that R2, which produces less fusel alcohols and isoacids throughout the fermentation, when compared with Opale might more effectively regulate carbon flux (less efficient usage of nitrogen) over time. Moreover, this aspect could be related to nitrogen causing the excretion of less carbon metabolic excess from the cells (40).

Contrary to VOC product formation, nutrient consumption behavior of nitrogen and sugar compounds is similar among the strains. It was observed that nitrogenous compound uptake began before the onset of growth in which ammonium was fully consumed first, similar to what was found in a previous study (41). There was a preference for specific amino acids noticed following ammonia consumption (42). These preferred amino acids were Asp, His, Glu, Met, and Lys, all consumed within 44 hours. These were also previously seen as initially consumed amino acids in media with the same YAN conditions (43, 44). This preference in utilization of particular amino acids is thought to be linked to nitrogen catabolite repression (NCR) of amino acid transport permeases (24, 45). Many of the amino acids were consumed during the first 96 hours of fermentation. However, some showed interesting kinetic behavior. Ala, Asn, and Glu showed a slight uptick in concentration during the stationary growth phase, which could be attributed to excretion before autolysis, as has been seen previously (46) though there are some differences regarding the type of amino acids.

The results from PLS linear regression modeling indicate that the production of VOCs is a function of nitrogen utilization. Previous studies have demonstrated that containing adequate YAN levels at the beginning of fermentation is essential in determining yeast cell growth and producing desirable levels of VOCs (8, 27, 47). The production of VOCs such as fusel alcohols and their associated acetate esters stems from the catabolism of branched-chain and aromatic amino acids via the Ehrlich pathway (48, 49). Furthermore, the availability of specific nitrogenous compounds early on during fermentation alters the transcriptional regulation of genes involved with higher alcohol production and the formation of esters in yeast (50, 51). Naturally, upon reaching the stationary growth phase, the rate of production of VOCs by *S. cerevisiae* begins to decline (29). It is understood that esters play a role in the survival of yeast in the esterification of toxic MCFAs, thus facilitating their diffusion through the plasma membrane (52).

By comparing the correlations of concentrations all other VOCs with nitrogen utilization (*R*^2^ ≥ 0.77) on the one hand and methionol concentration with nitrogen utilization (*R*^2^ = 0.61) on the other hand, it appears likely that nitrogen utilization is not the only significant factor influencing methionol production. The relatively low correlation for methionol formation could be due to the absence of causal relationships between the metabolic pathways of methionol biosynthesis and nitrogen metabolism. Methionol biosynthesis is also intricately linked to the sulfate reduction pathway and thus, the relationship of sulfur uptake during fermentation (53). Hence, this could explain the variation in PLS model predictions with other fusel alcohols. Previous works have confirmed that the role of central carbon metabolism in the synthesis of higher alcohols is significant akin to the contribution of amino acids (54). However, it is assumed that there is a balance between the amount of α-ketoacids supplied by central carbon metabolism and the amount of α-ketoacids converted into amino acids for protein biosynthesis. Thus, over the course of nitrogen utilization the corresponding flux of α-ketoacids related to the nitrogen utilization over time would be negligible compared to the flux originating from the central carbon metabolism (55). The α-ketoacid pool available would then remain unchanged and not affect the production of fusel alcohols.

The regression vector plots from both PLS regression models indicated numerous nitrogenous compounds in common among the models and the degree to which the nitrogen sources had influenced the models. The PLS models for total aroma production and biomass concentration indicated some shared nitrogenous compounds that are positively correlated with the formation of biomass and VOCs. These amino acids were glycine and lysine. Furthermore, for total aroma it was determined that the most significant positively correlated amino acids were glycine, lysine, leucine, and tyrosine. These findings corroborate what has been shown in a previous study, albeit with beer yeasts (32). This correlation is likely caused by the direct connection to the formation of some of the most significant fusel alcohols (isobutanol, isoamylol, and propanol) and their subsequent acetate esters (isobutyl acetate, isoamyl acetate, and propyl acetate) (56, 57).

pFBA was performed to determine the essential and metabolically feasible distribution of fluxes throughout the yeast metabolic network. By employing pFBA and FEA, the significance among essential reactions was observed for nitrogenous utilization pathways as well as sterol, pyruvate, and glycerolipid metabolism. A similar approach has been taken previously with *Escherichia coli* to understand network effects of iron metabolism in triggering oxidative stress in *Caenorhabditis elegans* (58)For instance, our analysis illustrates the link lysine plays in the formation of VOCs. In particular, this link is implied from the modeled essentiality among fluxes through amino acid permeases mainly LYP1 regulated permeases which have been suggested to steer higher alcohol and acetate ester formation during beer fermentations (33). The pFBA revealed many essential reactions in leucine utilization pathways through the use of permeases, transaminases, and decarboxylases alluding to the Ehrlich pathway to produce VOCs (49). This result suggests as, shown by Yoshimoto and coworkers, that overexpressing genes such LEU and BAT genes within leucine metabolism can increase isoamyl alcohol and isoamyl acetate secretion.

In summary, the commercial yeast strains investigated in this study were selected based on their aroma-producing properties and their favorable fermentation attributes at similar nitrogen levels to compare differences in their aroma formation throughout anaerobic fermentation. By applying HS-SPME/GC-MS and UPLC, we were able to quantitatively analyze production of key VOCs and consumption of nitrogenous compounds by these various yeast strains over multiple time points at different phases of fermentation. This provided unique insight into the specific aroma production profiles of these strains and how they change over time and between strains. Partial least-squares regression analysis indicated a strong correlation from the nitrogen utilization with two essential wine fermentation criteria: biomass growth and VOC production in *S. cerevisiae*. The PLS regression models also pointed to several key amino acids known to be involved in forming VOCs that are desired in wines. The application of genome-scale metabolic modeling provided a further mechanistic link to support the observed PLS correlations. Although there are many possible explanations for why yeast produce VOCs, the results demonstrated in this work indicate that nitrogen sources such as leucine, lysine, glycine, and tyrosine are positively correlated with yeast cell growth and VOC production.

Further work measuring other VOCs, sulfur and lipid compounds used in those corresponding assimilation pathways could provide greater insight into aroma formation mechanisms. Furthermore, gene expression profiling of commercial yeast during the fermentation process would help identify and understand possible strain-dependent behavior. This coupled with metabolic models examining the gene, enzyme, metabolic flux relationships, aroma formation, and nitrogen utilization, as has been performed to limited degree in other conditions (59, 60), would provide insight into elucidating the metabolic sources of VOCs formation. Nevertheless, the correlations established in this work between nitrogen utilization and aroma production facilitates the direction of further studies.

## MATERIALS AND METHODS

### Yeast strains

All four yeast strains - Uvaferm 43^TM^(Uvaferm), Lalvin R2^TM^ (R2), Lalvin ICV Opale^TM^ (Opale), Vitilevure^TM^ Elixir Yseo (Elixir), used in this study are Lallemand (Lallemand, Montreal, Quebec) commercial yeast strains. Uvaferm and R2 are*. Saccharomyces bayanus* which is a hybrid of *Saccharomyces cerevisiae*, *Saccharomyces eubayanus* and *Saccharomyces uvarum*: while Opale and Elixir are *S. cerevisiae* var. *cerevisiae*. All yeast strains were obtained from the UC Davis Enology Culture Collection. Additionally, yeast strains were chosen to have different fermentation and aroma-producing characteristics under the conditions selected. For instance, R2 and Elixir are reported to be wine yeast strains that produce relatively high amounts of esters. In contrast, Uvaferm is reported to be more of an aroma-neutral strain and Opale is reported to impart an enhanced, complex character to wines (Lallemand, Montreal, Quebec). The strains were stored at −80 °C in a 25% (v/v) glycerol solution according to the method of Amberg et al. (61). The strains were streaked for single colony isolation on yeast extract peptone dextrose (YEPD) agar plates, incubated at 25 °C for 24 - 48 h until sufficient colonies formed, and stored at 4 °C for no longer than 30 days.

### Growth, fermentation media, and culture conditions

MMM synthetic grape juice (220 g sugar/liter; 22.0 °Brix), with a 1:1 mixture of glucose (110 g/liter) and fructose (110 g/liter), 123 mg/L YAN, and 11 mg/L ammonium, was prepared according to the method of Giudici and Kunkee (62). MMM medium was used within 24 hours of preparation. Fermentations were carried out in 500 mL Erlenmeyer flasks sealed with a rubber stopper and air lock with a working volume of 400 mL similarly as described in Henderson et al. (63). In addition. the fermentations were inoculated according to the method stated in Henderson et al. (63) and likewise a volume of the inoculum to give an initial OD_600_ of ∼ 0.1 (∼15ml) was added to 400 ml of MMM medium. The pH of the MMM medium was 3.25, and the fermentation temperature was maintained at 20 °C. Cultures were stagnantly cultivated at 20 °C for 17 days. Initial cell concentration (OD_600_) and °Brix measurements were performed at the beginning of fermentation and subsequently ten more times over the course of the fermentation until °Brix fell below one or remained constant between two consecutive measurements. °Brix measurements were performed with a refractometer, and OD_600_ measurements were performed with a spectrophotometer. 10 mL samples were taken at regular intervals and transferred to a 15 mL Greiner tube, sealed, and frozen at −20 °C until analysis. Experiments were performed as biological triplicates.

### Cell dry weight determination

The total cell dry weight (biomass) of the samples was determined by taking a sample directly from the culture medium and passing through dried and pre-weighed membrane filters with a pore size of 0.2 μm (Pall Corporation, Ann Arbor, MI, USA) by a vacuum filtration unit as described by van Mastrigt and co-workers (64). Total biomass concentration at every sample time point was determined by transforming results from OD_600_ measurements using a calibration curve for each yeast strain. Total cell dry weights were determined in triplicates.

### HPLC analysis

After taking a sample from the Erlenmeyer flask, cells were immediately removed by centrifugation (13,000 x g for 10 min at 4°C), and the supernatant was stored at −20°C until analysis. Supernatants were deproteinated by adding 0.25 ml Carrez A (0.1 M potassium ferrocyanide trihydrate) and 0.25 ml Carrez B (0.2M zinc sulfate heptahydrate) to 0.5 ml sample followed by centrifugation for 10 min at 13000 x g. Glucose, fructose, lactate, acetate, malate, citrate, ethanol, 1-propanol, and 1,2-propanediol were quantified by High Performance Liquid Chromatography (HPLC) on a Ultimate 3000 (Dionex, Idstein, Germany) equipped with an Aminex HPX-87H column (300 x 7.8 mm) with precolumn (Biorad) as described by van Mastrigt and co-workers (64). As the mobile phase, 5 mM sulfuric acid was used at 0.6 ml min^-1^, and the column was maintained at 40°C. The injection volume was 10 µl. Compounds were identified by a refractive index detector (RefractoMax 520) for quantification and UV measurements at 220, 250, and 280 nm for identification. All analysis was performed in duplicate.

### UPLC amino acid analysis

40 μL of 5-fold diluted samples was mixed with 50 μL of 0.1 M HCl solution containing 250 μM norvaline as internal standard. Then, 10 μL of chilled 30% sulfosalicylic acid (SSA) was added, and the solution was mixed and centrifuged (13,000 x g) for 10 min. at 4 °C. Amino acids were derivatized using the AccQ•Tag Ultra derivatizaton kit (Waters). 20 μL of the deproteinated sample solution supernatant or standard amino acids mixture was mixed with 60 μL of a modified AccQ•Tag Ultra borate buffer (for deproteinated samples 150 μL of 4 M NaOH was added to 5 mL borate buffer). Next, 20 μL of a AccQ•Tag reagent previously dissolved in 2.0 mL AccQ•Tag Ultra reagent diluent was added and vortexed for 10 seconds. Then, the sample solution was capped and warmed at 55°C in a heatblock for 10 min. Amino acids and ammonium were quantified by Ultra-Performance Liquid Chromatography (UPLC) on an Ultimate 3000 (Dionex, Idstein, Germany) equipped with a AccQ•Tag Ultra BEH C18 column (150 mm x 2.1 mm, 1.7 µm) (Waters, Milford, MA, USA) with BEH C18 guard column (5 mm x 2.1 mm, 1.7 µm) (Waters, Milford, MA, USA). The column temperature was set at 55°C, and the mobile phase flow rate was maintained at 0.7 mL/min. Eluent A was 5% AccQ•Tag Ultra concentrate solvent A and Eluent B was 100% AccQ•Tag Ultra solvent B. The separation gradient was 0-0.04 min 99.9% A, 5.24 min 90.9% A, 7.24 min 78.8% A, 8.54 min 57.8% A, 8.55-10.14 min 10% A, 10.23-17 min 99.9% A. One microliter of sample was injected for analysis. Compounds were detected by UV measurement at 260 nm.

The ammonium concentration was further determined and confirmed with an ammonia assay kit (Megazyme, Bray, Ireland) according to the manufacturer’s procedures. The ammonia levels were verified to exhaust after 28 hours of fermentation. This method was also used to correct quantification of ammonium detected from the UPLC. Norvaline was used as an internal standard (IS) for the UPLC amino acid analysis. All analysis was performed with a single replicate.

### Volatile organic compounds (VOC) analysis

To determine the volatile organic compounds (VOCs), 2 mL sample was transferred to a 5 mL GC vial. Samples were stored frozen (−20 °C) until analysis by headspace solid-phase microextraction gas chromatography mass spectrometry (HS SPME GC-MS) (65). Samples were thawed and incubated for 5 min at 60 °C with agitation. Subsequently, VOCs were extracted from the samples for 20 min at 60 °C using a Solid-Phase Microextraction fiber (50μm Bonded Gray Hub (DVB/CAR/PDMS) Supelco, USA). The compounds were desorbed from the fiber for 10 min on a Stabilwax®-DA Crossband®-Carbowax®-polyethylene-glycol column (30m length, 0.25 mmID, 0.5 mm df). The settings on the gas chromatograph were PTV Split-less mode 5 min at 250 °C. Helium was used as carrier gas at a constant flow of 1.5mL/min. The temperature of the GC oven was initially at 40 °C. After 2 min, the temperature was raised to 240 °C at a rate of 10 °C/min and kept at 240 °C for 5 min. Mass spectral data was collected over a range of m/z 33-250 in full scan mode with 3.0030 scans/second. VOC profiles were analyzed with Chromeleon™ 7.3 software. The ICIS algorithm was used for peak integration and the NIST main library was used for identification by matching mass spectral profiles with the profiles in NIST. One quantifying peak (in general the highest m/z peak per compound) was used per compound for quantification, while one or two confirming peaks were used for confirmation.

The 11 aroma associated compounds studied were propan-1-ol, 3-methylbutan-1-ol (isoamylol), 2-methylpropan-1-ol (isobutanol), 2-phenylethan-1-ol, methionol, ethyl ethanoate (ethyl acetate), ethyl butanoate, ethyl hexanoate, 3-methylbutyl acetate (isoamyl acetate), 2-methylpropyl ethanoate (isobutyl acetate), and 2-phenylethyl acetate (Table 4). All of the purchased analytes had a purity of at least 98% and were purchased from Sigma-Aldrich (Sigma-Aldrich, Germany). Higher alcohols and esters were quantified using standard solutions in unfermented synthetic grape juice as similarly as described in the literature (29, 66). The model solutions were spiked with varying amounts of the 10 aroma compounds (higher alcohols and esters) to yield concentrations within the range of the concentrations typically found in unwooded and unaged wines (Table 4). The headspace was sampled by SPME GC-MS in the same manner as used for the wine samples. Standard curves of each studied aroma compound were created by plotting each aroma compound’s peak area against the standard concentration from the standard solutions. All samples were analyzed in duplicate.

**TABLE 4.**
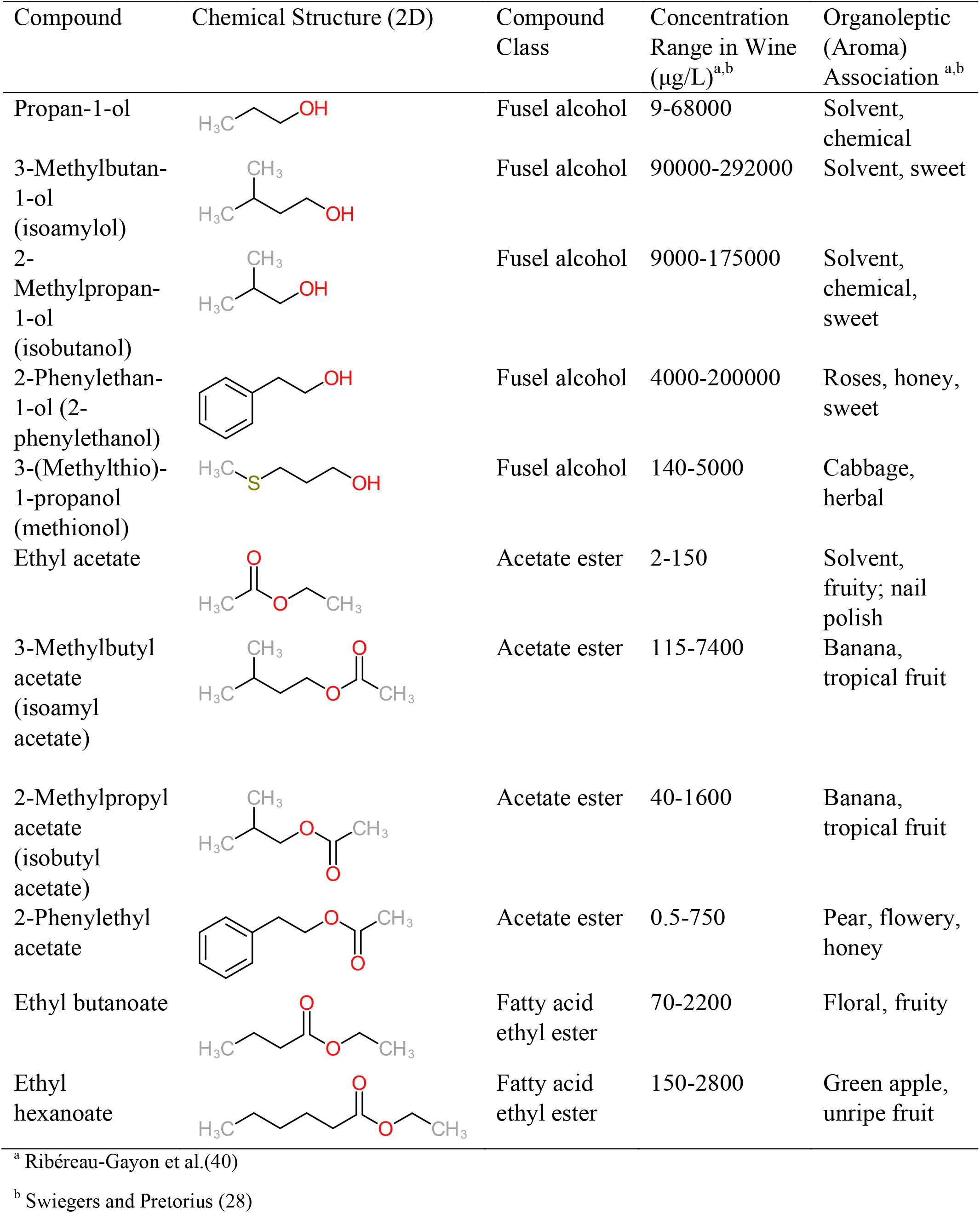
Summary of major of VOCs found in wines along with their chemical attributes

| Compound | Chemical Structure (2D) | Compound Class | Concentration Range in Wine (µg/L)<sup>a,b</sup> | Organoleptic (Aroma) Association<sup>a,b</sup> |
| --- | --- | --- | --- | --- |
| Propan-1-ol |  | Fusel alcohol | 9-68000 | Solvent, chemical |
| 3-Methylbutan-1-ol (isoamylol) |  | Fusel alcohol | 90000-292000 | Solvent, sweet |
| 2-Methylpropan-1-ol (isobutanol) |  | Fusel alcohol | 9000-175000 | Solvent, chemical, sweet |
| 2-Phenylethan-1-ol (2-phenylethanol) |  | Fusel alcohol | 4000-200000 | Roses, honey, sweet |
| 3-(Methylthio)-1-propanol (methionol) |  | Fusel alcohol | 140-5000 | Cabbage, herbal |
| Ethyl acetate |  | Acetate ester | 2-150 | Solvent, fruity; nail polish |
| 3-Methylbutyl acetate (isoamyl acetate) |  | Acetate ester | 115-7400 | Banana, tropical fruit |
| 2-Methylpropyl acetate (isobutyl acetate) |  | Acetate ester | 40-1600 | Banana, tropical fruit |
| 2-Phenylethyl acetate |  | Acetate ester | 0.5-750 | Pear, flowery, honey |
| Ethyl butanoate |  | Fatty acid ethyl ester | 70-2200 | Floral, fruity |
| Ethyl hexanoate |  | Fatty acid ethyl ester | 150-2800 | Green apple, unripe fruit |

<sup>a</sup> Ribéreau-Gayon et al.(40)
<sup>b</sup> Swiegers and Pretorius (28)

### Multivariate data and statistical analyses

The yeast strain-specific impact on the kinetics of amino acid consumption and aroma compound formation for the 11 volatile aromas were determined by plotting the compound concentrations throughout the fermentation. Furthermore, several performance metrics were accessed from the dynamic consumption and production profiles, such as the final concentration of volatile aroma compounds and utilization of nitrogenous compounds. In this study, nitrogen (ammonia/amino acids) utilization was determined as the difference in the ammonia/amino acid concentration from beginning to sample time point of interest throughout the fermentation (29). The final volatile aroma concentration was determined at 404 hr when all of the fermentations were finished. Principal component analysis (PCA) was performed using R (version 3.6.2, R Core Team, 2020).

One-way analysis of variance (ANOVA) (significance level of p < 0.05) was performed in R (version 3.6.2) using the statistics package applying post hoc Tukey’s HSD test (significance level of p < 0.05) as well as in SPSS (IBM SPSS Statistics 27, IBM Corp, Armonk, NY).

A multivariate statistical method known as partial least squares (PLS) regression was employed to determine a relationship between biomass and VOC production data (Y-Block) and nitrogen (ammonia and amino acid) utilization data (X-Block). The biomass and VOC production are defined as the difference of biomass and VOC concentration at the end of fermentation to at a sampled time point. Conversely, utilization was calculated as the difference in concentration at a sampled time point to end of fermentation. These two data blocks were imported into MATLAB (version MATLAB® (2017b), MathWorks, Natick, MA) for PLS regression analysis and interval-PLS (iPLS) variable selection using the PLS Toolbox (version 8.9; Eigenvector Research, Inc., Manson, WA). Parameters for iPLS were selected according to the method of Wise et al. (67) and Anderson and Bro (68) and adapted from the technique of Henderson et al. (63). Reverse-analysis-mode iPLS selection was conducted with an interval size of one variable with a maximum of 18 latent variables (LVs). The step size, which is the space between interval centers, and number of variables chosen were automatically decided based on when there was no improvement in the root mean squared error in cross-validation (RMSECV).

Subsequently, once iPLS was completed, the selected variables were added to PLS Toolbox for structured equation modeling. The used approach was adapted based on an approach by Henderson et al. (63) using the SIMPLS algorithm (69) at a confidence limit of 0.95. Auto-scaling followed by mean centering was applied to the preprocessing of the X- and Y-blocks. The PLS model was cross validated using Venetian Blinds with 24 data splits and a maximum of 18 LVs. The quality and predictive capability of the PLS model were evaluated from the correlation coefficient (*R*^2^), cross-validated correlation coefficient (Q^2^), root mean squared error of correlation (RSMEC), and RMSECV. The impact of selected nutrient utilization variables on the PLS model was assessed using scores, loadings, and regression vector plots. Overall, the PLS model was used to determine the nutrient utilization (ammonia and amino acids), which contribute to the biomass formation and VOCs production during fermentation.

### Metabolic modeling

For model analysis, we used the latest genome-scale metabolic model (GSMM) of *S. cerevisiae*, Yeast 8.4.2 (70). To adequately model an anaerobic fermentation, we proceeded as suggested by Heavner et al. (71), constraining *v_o_2__* to zero (LB=UB=0 [mmol/g DW h]), allowing unrestricted uptake of ergosterol (r_1757), lanosterol (r_1915), zymosterol (r_2106), 14-demethyllanosterol (r_2134), and ergosta-5,7,22,24(28)-tetraen-3beta-ol (r_2137) and oleate (r_2189). In addition, pathways including the oxaloacetate-malate shuttle and glycerol dehydrogenase reaction were unrestricted as described by Sanchez et al. (72, 73) (in the model this was achieved by blocking reactions r_0713, r_0714 and r_0487). Heme A was also removed from the biomass equation as it is not used under anaerobic conditions. Experimentally-derived net uptake and production fluxes which were taken from measured data (Fig. 1, Fig. 2, and Fig. 3) were used to constrain the model in the form of exchange reactions (LB=UB) for the sugars, amino acids, organic acids, VOCs and other byproducts.

We applied parsimonious flux balance analysis (pFBA) to evaluate which reactions and pathways are essential under the given system conditions and eliminate blocked reactions. It uses a bilevel optimization in which the growth rate (biomass) is optimized using FBA, followed by the minimization of total flux through all gene-associated reactions. The underlying assumption is that, under growth pressure, there is a selection for strains that can reach highest growth yield while using the minimum amount of enzyme. Then, genes / reactions are classified into six categories based on Lewis et al. (74), i.e. 1) essential genes, metabolic genes necessary for growth in the given media; 2) pFBA optima, non-essential genes contributing to the optimal growth rate and minimum gene-associated flux; 3) enzymatically less efficient (ELE), genes requiring more flux through enzymatic steps than alternative pathways that meet the same predicted growth rate; 4) metabolically less efficient (MLE), genes requiring a growth rate reduction if used; 5) pFBA no-flux, genes that are unable to carry flux in the experimental conditions; and 6) blocked, genes that are only associated with the reactions that cannot carry a flux under any condition (“blocked” reactions). Flux enrichment analysis (FEA) was applied to evaluate enrichment by comparing the determined frequency of an annotated pathway with the frequency expected by chance within the model. The statistical significance for the pathways or reaction subsystems to be enriched contained p < 0.01. All reactions, metabolites and subsystems are defined according to the Biochemical Genetic and Genomic (BiGG) database convention (75).

All metabolic modeling was performed in MATLAB (version MATLAB® (2017b), MathWorks, Natick, MA) using the using Cobra Toolbox 3.0 (76).

## ACKNOWLEDGEMENTS

This project was supported by the American Vineyard Foundation grant number 2252 and the Ernest Gallo Endowed Chair in Viticulture and Enology. In addition, materials and consumables were funded by a Wageningen University and Research internal fund.

We thank Judith Wolkers–Rooijackers for her assistance with the sampling and experimental analysis. In addition, we also appreciate Lucy Joseph for helping us obtain the wine yeast strains from the University of California Davis culture collection.

